# Extracellular proteomic profiling of *Bacillus toyonensis* biovar Thuringiensis Bto_UNVM-42 identifies candidate proteins potentially associated with its nematicidal activity

**DOI:** 10.64898/2026.04.06.716753

**Authors:** Sergio Redondo-Moreno, Cecilia Peralta, Leopoldo Palma

## Abstract

Previous studies demonstrated that cell-free supernatants of *Bacillus toyonensis* biovar Thuringiensis Bto_UNVM-42 exhibit nematicidal activity against *Panagrellus redivivus*, although the molecular basis of this phenotype remains unclear. While pesticidal proteins in *Bacillus thuringiensis* and related species are classically considered intracellular and associated with parasporal crystals, their occurrence in extracellular fractions remains poorly explored. Here, we characterized the extracellular protein repertoire of biologically active supernatants and evaluated their effects on *Caenorhabditis elegans* N2. LC–MS/MS analysis of LB-derived cell-free supernatants revealed the presence of pesticidal protein homologs related to Cry32-, Cyt1, and Mpp3-like protein families, together with degradative enzymes including collagenases, chitinases, proteases, phospholipases, and cytolysins. Signal peptide prediction suggested classical secretion pathways for several proteins, whereas others may reach extracellular fractions through alternative localization mechanisms. Comparative DIA-based proteomic analyses of LB and CCY supernatants revealed marked medium-dependent differences in the extracellular protein repertoire, including differential enrichment of Bt-like pesticidal homologs and degradative enzymes associated with distinct physiological states. Exposure assays showed that concentrated cell-free supernatants from both conditions caused severe loss of mobility and morphological alterations consistent with tissue disruption in *C. elegans*. Together, these findings provide proteomic evidence for the extracellular occurrence of pesticidal and degradative protein homologs in Bto_UNVM-42 and identify candidate extracellular factors potentially associated with the nematicidal activity of this strain. More broadly, the results support the view that soluble extracellular components may contribute to the biological activity of Bt-related bacteria beyond the classical spore–crystal paradigm.

## 1. Introduction

*Bacillus thuringiensis* and related members of the *Bacillus cereus* group are widely recognized for their production of pesticidal proteins, particularly Cry and Cyt proteins, which have been extensively exploited in biological control (Slamti and Lereclus, 2026). However, growing evidence indicates that factors beyond classical crystal proteins may also contribute to pathogenicity, including extracellular enzymes and accessory proteins involved in host interaction and tissue degradation (Jouzani et al., 2017; Niu et al., 2007).

Previous studies have contributed to expanding the known ecological and pesticidal diversity of *B. toyonensis* biovar Thuringiensis. In 2022, we reported the first *B. toyonensis* biovar Thuringiensis strain (NCBI Taxonomy ID: 2923195) exhibiting dual insecticidal activity against both lepidopteran and coleopteran pests (Sauka et al., 2022). More recently, a comprehensive genomic and biological characterization highlighted the overlooked entomopathogenic potential of this taxon and its relationship to classical Bt-like bacteria (Sauka et al., 2026a). In addition, strain Bto_UNVM-42 was shown to display nematicidal and nematostatic activity against *Panagrellus redivivus* and *Pratylenchus zeae* (Sauka et al., 2026b), respectively, raising questions regarding the molecular basis of these phenotypes. Genome analysis of this strain (Acc. No. JAYEPS000000000) revealed a diverse repertoire of putative pesticidal genes encoding for Cry32-, Cyt1-, and Mpp3 Bt-like homologs, together with several genes encoding proteins potentially associated with host interaction and tissue degradation, such as chitinases, enhancin-like proteins, and Sep1-like serine proteases (peptidase S8 superfamily) (Fang et al., 2009; Geng et al., 2016).

Although genomic analyses provide valuable insights into the pathogenic potential of bacterial isolates, they do not reveal which proteins are actually present in biologically active extracellular fractions. Consequently, the composition of the extracellular protein repertoire of Bto_UNVM-42 and its potential relationship with the previously observed nematicidal phenotype remain largely unexplored.

Since previous work demonstrated that the nematicidal activity of strain Bto_UNVM-42 was associated with cell-free culture supernatants rather than with bacterial cells or spores (Sauka et al., 2026b), the present study aimed to explore the extracellular protein repertoire potentially underlying this phenotype. We first performed an exploratory LC–MS/MS characterization of biologically active LB-derived cell-free supernatants, followed by a comparative proteomic analysis of extracellular fractions obtained from LB and CCY media, two culture conditions commonly used for vegetative growth and sporulation-associated protein production in *Bacillus* spp. In parallel, the biological effects of concentrated extracellular fractions were evaluated in *Caenorhabditis elegans* N2 as a tractable nematode model.

To our knowledge, reports describing the extracellular occurrence of Cry32-, Cyt1-, and Mpp3-like homologs remain limited, particularly in the context of comparative extracellular proteomic analyses. The detection of these proteins in biologically active cell-free supernatants expands current knowledge of Bt-related protein localization and provides new insights into the diversity of extracellular proteins produced by members of the *B. cereus* group.

## 2. Materials and methods

### 2.1 Culture media and supernatants preparation

*Bacillus toyonensis* biovar *thuringiensis* Bto_UNVM-42, previously described by Sauka et al. (2026), was grown in Luria-Bertani (LB) and CCY (Casein hydrolysate–Casamino acids–Yeast extract) media (Stewart et al., 1981) for 48 h at 28 °C with agitation (150 rpm). Following incubation, cultures were centrifuged at 13,000 × g for 10 min at 4 °C to remove bacterial cells and debris. The resulting supernatants were recovered and filtered through sterile 0.22 µm pore-size membranes to ensure the removal of residual cells. Cell-free supernatants were subsequently concentrated 10-fold using Amicon ultrafiltration units with a 10 kDa molecular weight cutoff (Millipore). Total extracellular protein concentration was determined using the Bradford assay (Bradford, 1976).

For the exploratory proteomic characterization, a single extracellular protein preparation obtained from LB-grown cultures was analyzed by LC–MS/MS to identify candidate extracellular proteins. For comparative proteomic analyses, three independent biological replicates were generated for each culture condition (LB and CCY) and analyzed using a DIA-based workflow. Bacterial cultures were monitored by phase-contrast microscopy (Nikon Eclipse) to assess cell morphology and physiological state prior to protein extraction.

### 2.2 Qualitative identification of extracellular proteins by DDA LC–MS/MS

LC–MS/MS analysis were performed at the Proteomics Service of the University of Valencia (Spain) using an Evosep One nanoLC system coupled to a timsTOF fleX mass spectrometer (Bruker).

Raw spectra were searched against a custom database containing the predicted proteome of strain Bto_UNVM-42 obtained from RAST annotation (Aziz et al., 2008), together with common contaminants and decoy sequences, using FragPipe v23.1 and MSFragger v4.3. Protein identifiers (e.g., peg.6278) correspond to protein-coding sequences predicted during RAST genome annotation. Peptide-spectrum matches, peptides, and proteins were filtered to a false discovery rate (FDR) of 1% using a target-decoy strategy. For the qualitative characterization of the extracellular proteome, only proteins supported by at least five unique peptides were retained for downstream analyses. Sequence coverage, peptide counts, and spectral abundance metrics were used exclusively for qualitative and semi-quantitative protein characterization and were not used for the subsequent DIA-based differential abundance analysis.

### 2.3 Bioinformatic analysis of extracellular proteins

Protein sequences were queried using BLASTp (Altschul et al., 1990) against a custom database derived from the Bacterial Pesticidal Protein Resource Center (BPPRC) (Berry et al., 2025), manually curated to include additional entries such as Sep1-like serine proteases (peptidase S8 family) and Bel enhancin homolog (Fang et al., 2009). Top-scoring hits were further validated using BLASTp searches against the NCBI non-redundant database, with particular attention to matches with proteins of known structure in the Protein Data Bank (PDB). For proteins classified as putative Bt-like pesticidal homologs (Cry32-, Cyt1-, and Mpp3-like), conserved domains were identified using Pfam through the InterProScan platform integrated in Geneious Prime v2026.1.0 (https://www.geneious.com).

Identified proteins were further analyzed for secretion signals using SignalP 6.0 (Teufel et al., 2022) and SecretomeP (Bendtsen et al., 2005) to predict classical and non-classical secretion pathways, respectively. In addition, Phobius was used to complement signal peptide prediction and to assess transmembrane features through the InterProScan platform integrated in Geneious Prime v2026.1.0.

To further characterize the functional composition of the extracellular proteome, proteins identified with at least five unique peptides were additionally subjected to BLASTp analysis against the UniProtKB/Swiss-Prot database (Boutet et al., 2007). Searches were performed locally using BLAST+ with an E-value threshold of 1e-^10^, retaining the top-scoring hit for each protein. Functional categories were assigned based on keyword-based annotation of Swiss-Prot descriptions, grouping proteins into degradative enzymes, pesticidal homologs, metabolic proteins, transport-associated proteins, and hypothetical/uncharacterized proteins. Only proteins with significant matches in Swiss-Prot were included in this classification analysis.

### 2.4 Comparative DIA-based proteomic analysis

A separate DIA-based proteomic experiment was performed to compare extracellular protein abundance between LB and CCY culture supernatants. Three independent biological replicates were analyzed per condition. Proteins from filtered and concentrated culture supernatants were digested with trypsin and analyzed by LC–MS/MS at the Proteomics Service of the University of Valencia (Spain) using a timsTOF platform operated in diaPASEF mode. Raw DIA data were processed using FragPipe v23.1 and DIA-NN v1.8.2 beta 8. Peptide identification and quantification were performed against a predicted spectral library generated from a custom protein database containing the predicted proteome of strain Bto_UNVM-42 and relevant pesticidal protein sequences. Peptide precursors and protein groups were filtered at a false discovery rate (FDR) of 1%. Protein abundance values obtained from the DIA-NN protein-group abundance matrix were used for downstream statistical analyses, including principal component analysis (PCA), hierarchical clustering, correlation analyses, and differential abundance analyses. Protein abundance estimates were based on DIA-derived quantitative signal intensities and were independent of the peptide counts, sequence coverage, and spectral abundance metrics used for the qualitative DDA-based protein characterization.

### 2.5 Qualitative nematode exposure assay

*Caenorhabditis elegans* strain N2 was maintained in our laboratories on nematode growth medium (NGM) agar plates seeded with *Escherichia coli* OP50 at 20 °C. Unsynchronized mixed developmental-stage nematodes were harvested from plates using M9 buffer and collected by centrifugation at 1,300 × g.

Approximately 500 nematodes were counted using a nematode counting slide (Microscope Services & Sales, UK) and exposed to 500 µL of concentrated cell-free culture supernatants obtained from LB and CCY cultures of Bto_UNVM-42. Assays were performed at 20 °C without agitation for 24 h. Control treatments consisted of sterile LB and CCY media processed under identical conditions.

Following incubation, nematodes were gently washed with 500 µL of M9 buffer to remove residual supernatant components and examined under bright-field microscopy to assess recovery of mobility and morphological alterations associated with loss of viability. Observations were performed using a Nikon Eclipse Ci microscope equipped with a digital 4K full HD camera.

## 3. Results

### 3.1 Cell morphology under different growth conditions

Phase-contrast microscopy of cultures after 48 h revealed clear differences between growth conditions (Figure 1). Cells grown in LB medium were predominantly observed as vegetative bacilli, with no detectable sporulation or formation of parasporal inclusions (Figure 1A). In contrast, cultures grown in CCY medium displayed abundant refractile structures consistent with spores and parasporal crystal formation (Figure 1B). These observations confirm that CCY promotes a sporulation-associated physiological state, whereas LB maintains conditions more suitable for the analysis of the soluble extracellular fraction.

**Figure 1.**
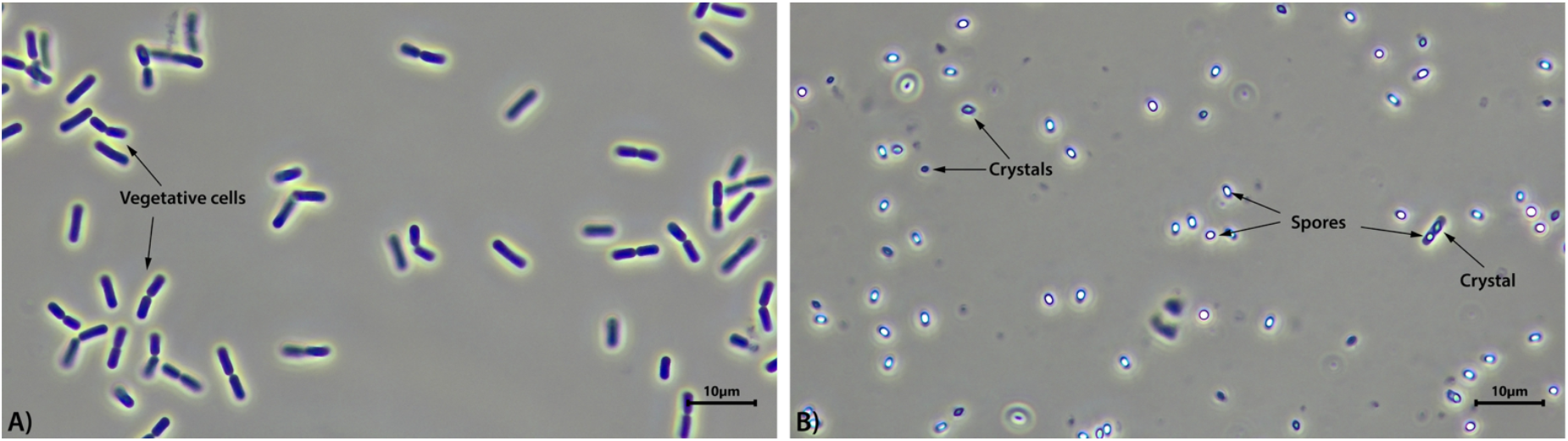
Phase-contrast microscopy of *B. toyonensis* Bto_UNVM-42 cultures after 48 h under different growth conditions. (A) LB-grown culture. (B) CCY-grown culture. Arrows indicate representative vegetative cells (A), spores and parasporal crystals (B). Images were acquired at 1000× magnification.

### 3.2 SDS–PAGE analysis of extracellular protein profiles

The extracellular protein profiles of culture supernatants obtained from LB and CCY media were analyzed by SDS–PAGE (Figure 2). Protein banding patterns were highly reproducible across the three independent biological replicates for each condition, indicating consistent sample preparation and protein recovery.

**Figure 2.**
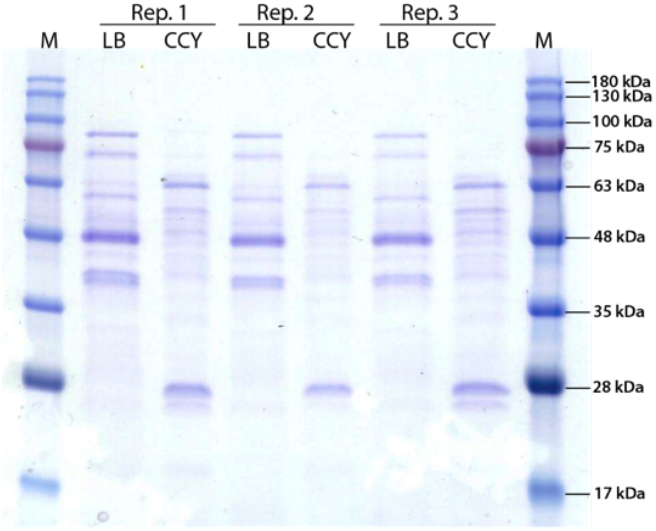
SDS–PAGE analysis of extracellular protein profiles from *B. toyonensis* Bto_UNVM-42.

Clear differences were observed between growth conditions. LB-derived supernatants displayed a more complex banding pattern, with multiple protein bands distributed across a broad molecular weight range, particularly between ~35 and 75 kDa. In contrast, CCY supernatants showed a comparatively simpler profile, with fewer detectable bands and lower overall signal intensity (Figure 2).

Bradford quantification of extracellular proteins revealed concentrations of 0.25, 0.21, and 0.37 mg/ml for LB biological replicates (LB1, LB2 and LB3), whereas CCY replicates (CCY1, CCY2 and CCY3) showed concentrations of 0.30, 0.31, and 0.40 mg/ml, respectively.

### 3.3 Qualitative identification of extracellular proteins by LC-MS/MS in LB medium

Given that nematicidal activity was previously detected in cell-free culture supernatants of strain Bto_UNVM-42 (Sauka et al., 2026b), we focused our analyses on extracellular fractions obtained after removal of bacterial cells. An initial exploratory LC–MS/MS analysis was performed using LB-derived supernatants, followed by comparative DIA-based proteomic analyses of LB and CCY extracellular fractions to evaluate the influence of growth conditions on the extracellular protein repertoire.

Among the 154 proteins detected in the LB culture supernatant, 96 were supported by at least five unique peptides and considered for downstream analysis. Of these, 53 showed significant similarity to curated Swiss-Prot entries (E-value ≤ 1e−10) and were assigned to functional categories (Figure 3).

**Figure 3.**
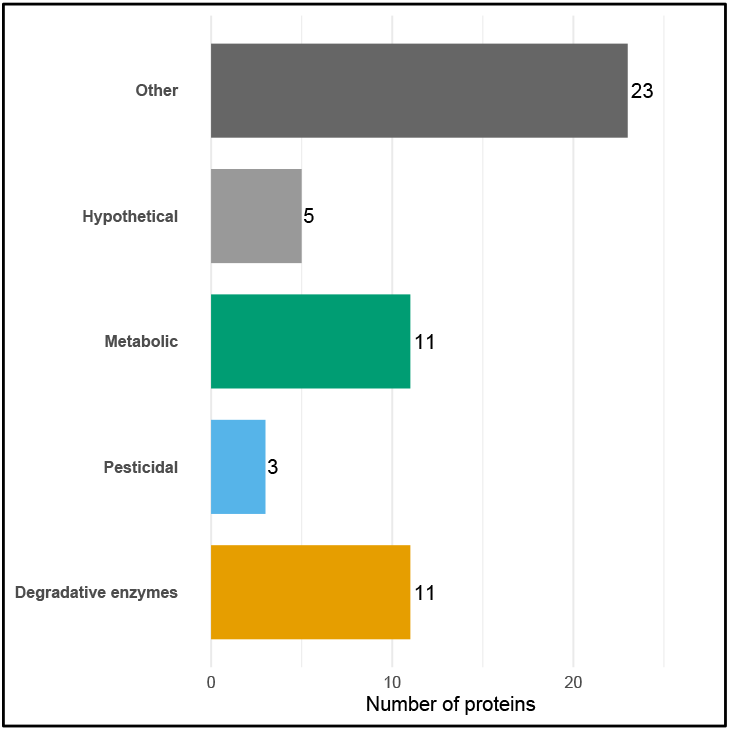
Functional classification of extracellular proteins identified in the culture supernatant of B. toyonensis Bto_UNVM-42. Proteins supported by at least five unique peptides and showing significant similarity to Swiss-Prot entries (E-value ≤ 1e−10) were grouped into functional categories based on annotation keywords.

Most of these proteins corresponded to degradative enzymes, predominantly proteases, together with collagenase and chitinase, followed by metabolic and transport-associated proteins. Notably, a smaller subset corresponded to pesticidal homologs, including Cry32-, Cyt1-, and Mpp3-like proteins. These proteins were identified in LB-derived supernatants obtained from cultures composed predominantly of vegetative cells and lacking detectable sporulation or parasporal crystal formation (Figure 1). Several additional proteins remained classified as hypothetical or uncharacterized. This distribution highlights the predominance of enzymatic components in the extracellular fraction, together with a defined but minor fraction of putative pesticidal proteins.

### 3.4 Bioinformatic analyses

Based on their homology to known pesticidal proteins and degradative enzymes, nine proteins were selected for further detailed analysis, including Cry32-, Cyt1-, and Mpp3-like proteins, as well as collagenase (ColA), cytolysins, chitinase, an S8 peptidase (Sep1-like), and a Bel enhancin homolog (Table 1). Among these, collagenase (ColA) and cytolysins exhibited the highest sequence coverage and peptide counts, whereas Cry32-like and enhancin-like proteins were detected with comparatively lower coverage.

**Table 1.**
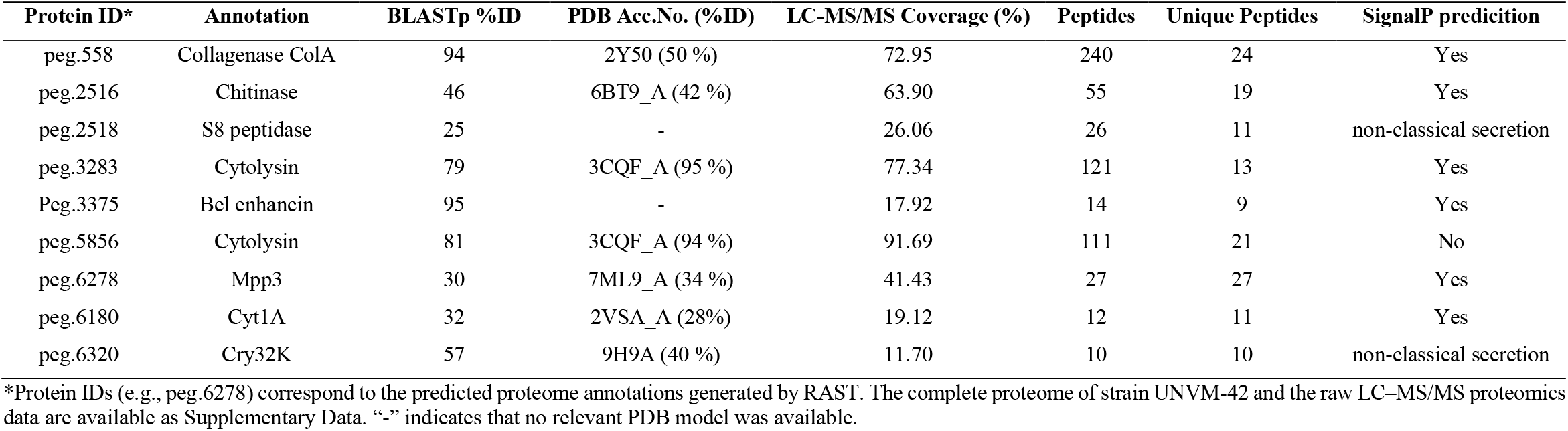
Proteins identified in the exploratory LC–MS/MS analysis of extracellular proteins recovered from LB culture supernatants of *B. toyonensis* strain Bto_UNVM-42. Signal peptide predictions were obtained using SignalP 6.0, and candidate non-classically secreted proteins were identified using SecretomeP. PDB accession numbers correspond to Protein Data Bank entries used for structural homology comparisons. Protein identifiers (e.g., peg.6278) correspond to gene models predicted from the genome annotation generated using RAST.

Notably, the cytolysin (peg.5856) was annotated by RAST as a truncated gene; however, its high coverage in the proteomic dataset supports that it is expressed and present in the extracellular fraction. This may reflect an automated annotation artifact or a genuinely shortened cytolysin-like variant that is nevertheless produced by the strain.

Signal peptide prediction using SignalP 6.0 supported classical secretion for most of the detected proteins, including the Mpp3-like protein, ColA, cytolysin, chitinase, Cyt1-like, and Bel enhancin homolog. In contrast, the Cry32 K-, G-, M-, and U-like homologs lacked predicted canonical signal peptides. Nevertheless, SecretomeP analyses supported their potential extracellular localization through non-classical secretion pathways, yielding positive SecP scores of 0.787, 0.916, 0.898, and 0.875, respectively. Similarly, the S8 peptidase did not present a predicted signal peptide but was identified as a candidate for non-classical secretion.

The Cry32-, Cyt1-, and Mpp3-like proteins showed conserved domain architectures consistent with known Bt-like pesticidal proteins (Figure 4 A, B and C). The Mpp3-like protein contained an ETX/MTX2 domain (InterPro ID: <u>IPR004991</u>), the Cyt1-like protein included a ricin B-lectin-like domain (InterPro ID: <u>IPR000772</u>), and the Cry32-like proteins displayed conserved protoxin-associated domains typical of Cry proteins, including Endotoxin_N (InterPro ID: <u>IPR005639</u>), Endotoxin_M (InterPro ID: <u>IPR001178</u>), Endotoxin_C (InterPro ID: <u>IPR005638</u>), Endotoxin_C2 (InterPro ID: <u>IPR054544</u>), Cry1Ac_D5 (InterPro ID: <u>IPR041587</u>), Cry1Ac_dom-VII (InterPro ID: <u>IPR048645</u>) and Crystall, a beta/gamma crystallin-like conserved domain region (InterPro ID: <u>IPR001064</u>).

**Figure 4.**
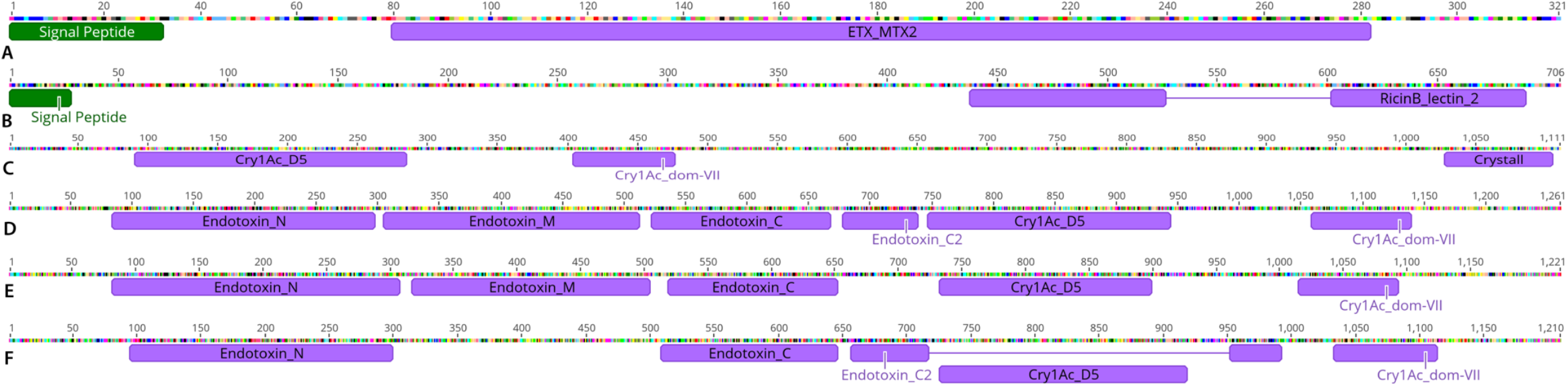
Conserved domain architecture of protein homologs identified in the proteomic profiling of *B. toyonensis* Bto_UNVM-42. (A) main organization of the Mpp3-like protein, displaying conserved motifs consistent with ETX/MTX2 family pesticidal proteins (Acc. No. PZ272960). (B) Domain architecture of the Cyt1-like protein, highlighting conserved cytolytic domains typical of β-sheet-rich Cyt toxins involved in membrane disruption (Acc. No. PZ272961). (C, D, E and F) Domain organization of the Cry32K, G, M and U-like proteins, respectively, showing the presence of conserved structural domains consistent with Cry protoxin-associated domains, including regions associated with pore formation and receptor binding (Acc. No. PZ272962). Cry32K, Cry32G, Cry32M and Cry32U homologs were identified as significantly enriched proteins in the comparative DIA-based analysis of LB and CCY extracellular proteomes. Green boxes indicate predicted signal peptides (Phobius), while purple boxes represent conserved domains identified using Pfam.

### 3.5 Comparative proteomic analysis of LB- and CCY-derived extracellular fractions

Comparative LC–MS/MS analyses revealed substantial differences in the extracellular proteomic profiles of *B. toyonensis* Bto_UNVM-42 grown in LB and CCY media. Differential abundance analysis identified numerous proteins that varied significantly between culture conditions, indicating marked changes in extracellular protein composition associated with each physiological state (Figure 5).

**Figure 5.**
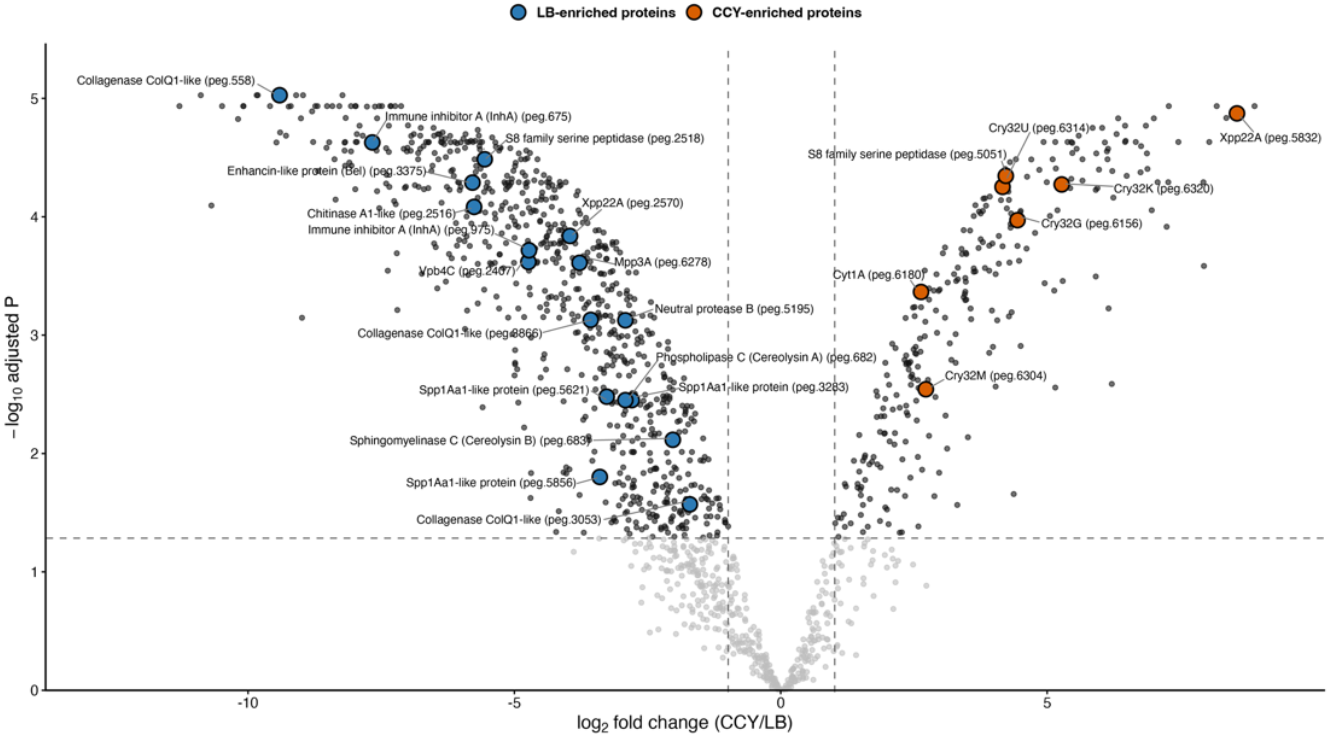
Volcano plot showing differential protein-group abundance between CCY and LB culture supernatants of *B. toyonensis* Bto_UNVM-42. The x-axis represents the log2 fold change for the CCY versus LB comparison, with protein groups enriched in LB shown on the left and those enriched in CCY shown on the right. Selected proteins discussed in the text are highlighted in blue when enriched in LB and in orange when enriched in CCY. Grey lines connect the highlighted points to their corresponding labels.

A total of 1,451 protein groups were detected across all samples, of which 1,011 showed significant differential abundance between LB and CCY conditions. Several proteins enriched in CCY supernatants corresponded to Bt-like pesticidal homologs, including Cry32-like proteins, Cyt1A and Vpb4C homologs, together with additional proteins associated with sporulation-related growth conditions (Table 2). In contrast, LB supernatants were enriched in a diverse repertoire of extracellular degradative enzymes and virulence-associated proteins, including collagenases, chitinases, immune inhibitor A (InhA)-like metalloproteases, phospholipase C, sphingomyelinase C, bacillolysin, neutral proteases, and enhancin-like proteins, as well as the pesticidal homolog Mpp3A-like (Table 2).

**Table 2.**
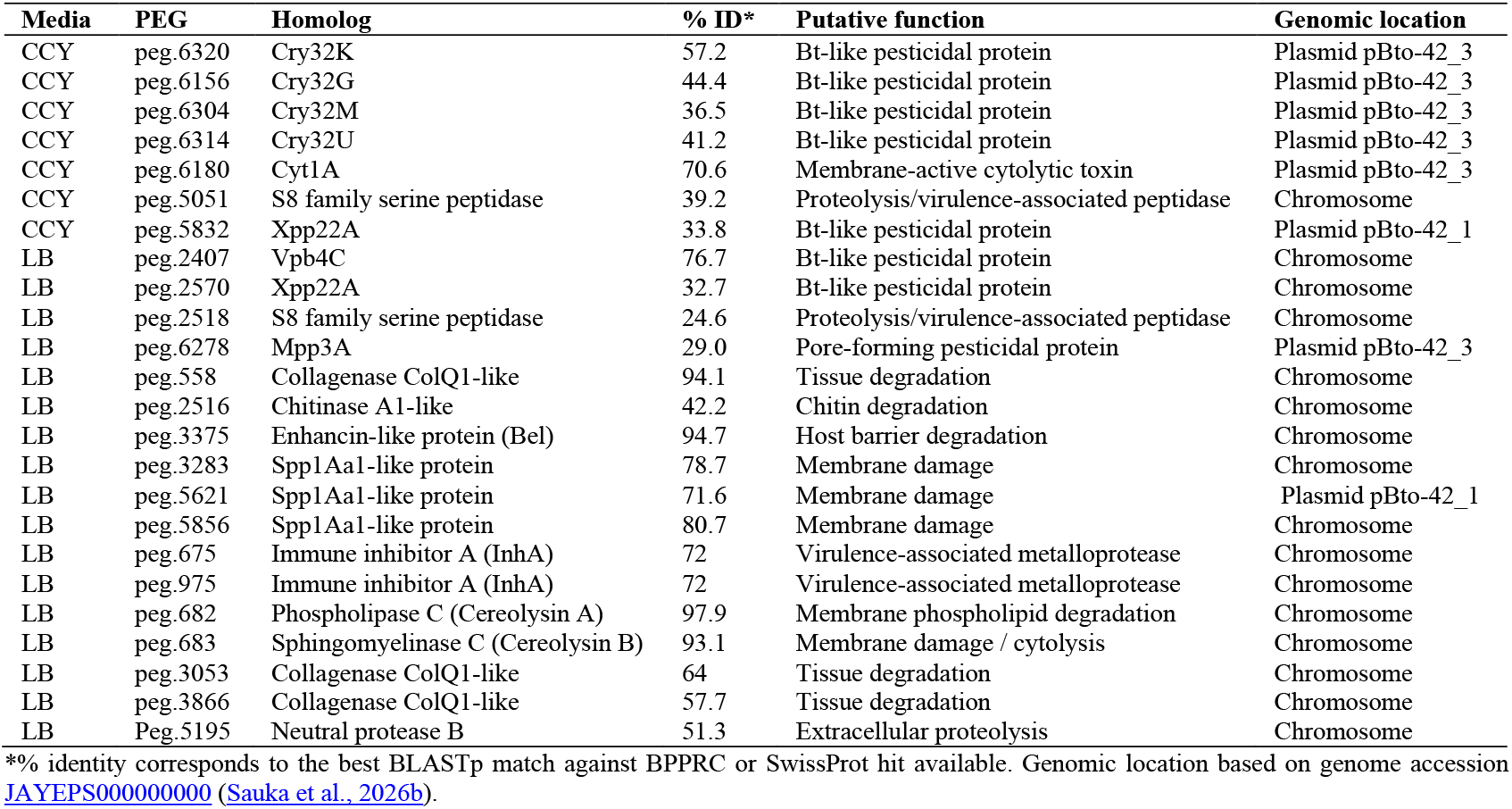
Selected extracellular proteins showing significant differential abundance between LB and CCY culture supernatants of strain Bto_UNVM-42. Proteins were selected based on differential proteomic analyses using an adjusted *p*-value cutoff ≤ 0.05 and |log_2_ fold change| ≥ 1.

All significantly differentially abundant proteins were detected in both LB and CCY cell-free supernatants. Consequently, differential abundance reflected quantitative changes in protein production between media rather than exclusive occurrence in a specific culture condition.

### 3.6 Effects of extracellular supernatants on nematode viability

After 24 h of exposure, nematodes treated with concentrated cell-free supernatants from both LB and CCY cultures showed complete absence of movement, whereas control nematodes maintained normal morphology and mobility. Importantly, nematodes did not recover mobility after washing with M9 buffer (Figure 6C and D). Although mortality was not formally quantified, all observed nematodes remained immobile after washing and exhibited morphological features consistent with death, suggesting that the treatments resulted in complete nematode mortality under the conditions tested.

**Figure 6.**
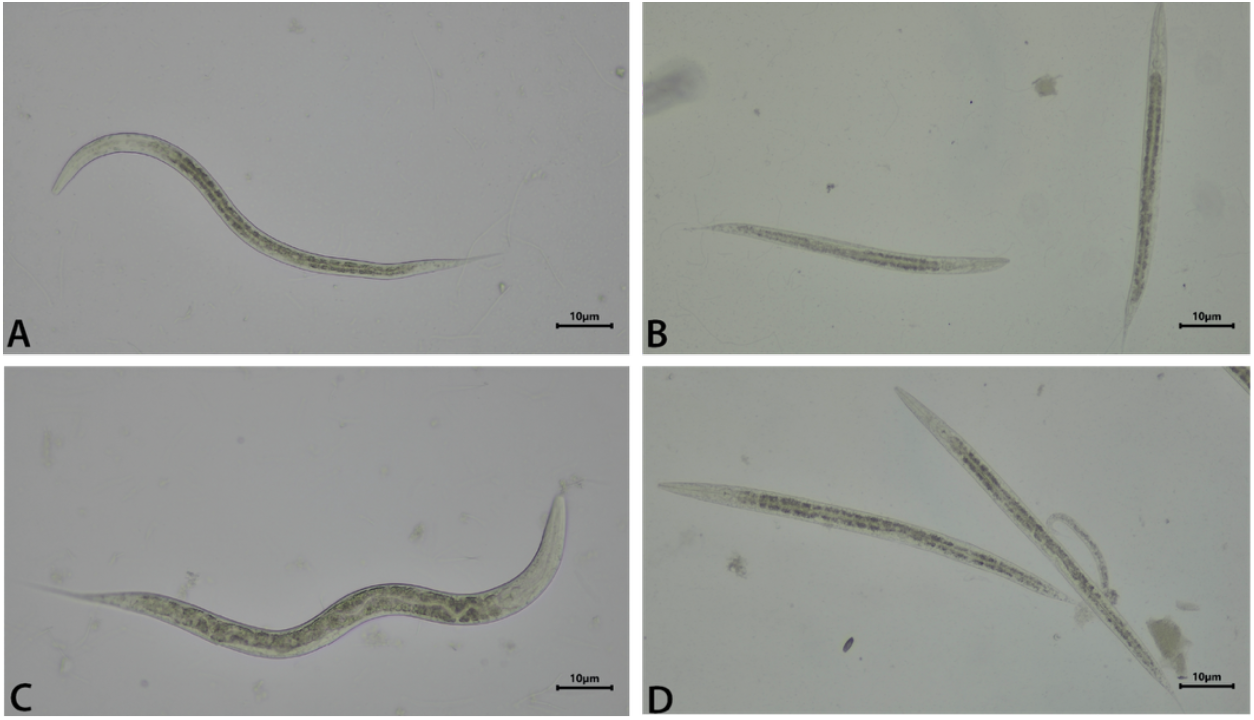
Effects of culture supernatants on *C. elegans* N2 after 24 h of exposure. Representative bright-field microscopy images of adult nematodes incubated with sterile media (controls) or cell-free concentrated supernatants. (A) Control in LB medium. (B) concentrated LB supernatant. (C) Control in CCY medium. (D) concentrated CCY supernatant.

In addition, some treated individuals displayed visible morphological alterations, including loss of internal structural definition and signs of tissue degradation (Figure 7 A, B, C and D), consistent with the pathological changes previously observed in *P. redivivus* exposed to the same cell-free supernatants (Sauka et al., 2026b).

**Figure 7.**
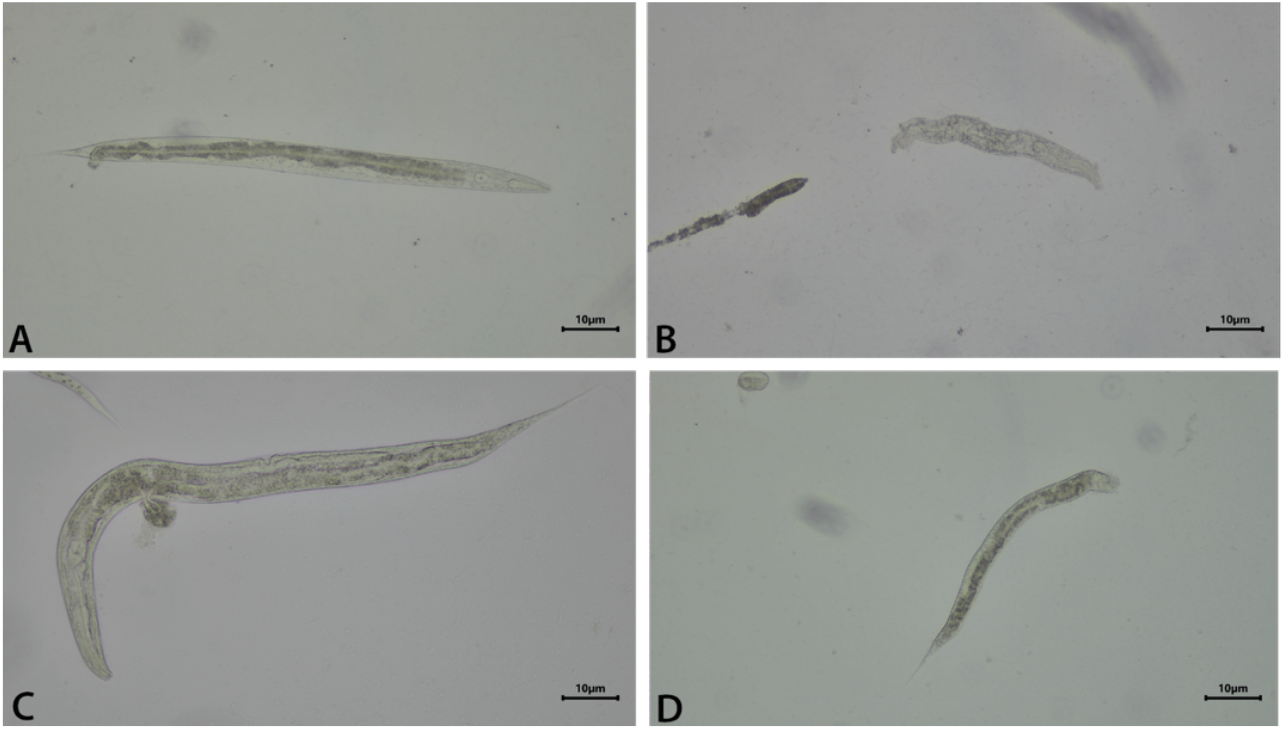
Morphological alterations, including swelling and tissue degradation in free-cell supernatants of LB (A and B) and CCY (C and D).

## 4. Discussion

A key finding of this study is the extracellular detection of Cry32-, Cyt1-, and Mpp3-like protein homologs in cell-free supernatants of *B. toyonensis* Bto_UNVM-42. Since the analyzed fractions were obtained from filtered culture supernatants, these results support the presence of soluble Bt-related proteins outside the classical spore–crystal complex. Although Cry- and Cyt-type proteins are generally regarded as intracellular components of parasporal inclusions released upon cell lysis, their detection in extracellular fractions in the present study suggests that alternative routes of release may occur under certain conditions (Schnepf et al., 1998; Slamti and Lereclus, 2026).

The comparative analyses performed under LB and CCY conditions further demonstrated that extracellular protein composition is strongly influenced by the physiological state of the culture. Phase-contrast microscopy confirmed that CCY promoted sporulation and parasporal inclusion formation, whereas LB cultures remained predominantly vegetative. These differences were consistent with the extracellular protein profiles observed by SDS-PAGE and DIA-based comparative proteomics, which revealed distinct proteomic compositions between both culture conditions. In particular, CCY supernatants showed a relative enrichment of Cry32- and Cyt1-like homologs together with sporulation-associated proteins, whereas LB supernatants displayed higher relative abundance of extracellular degradative enzymes, including collagenases, chitinases, phospholipases, metalloproteases, cytolysins, and enhancin-like proteins. These findings suggest that the extracellular protein composition of Bto_UNVM-42 may vary according to culture conditions.

An additional observation emerging from the comparative analyses was the apparent genomic partitioning of extracellular functions. Most Bt-like pesticidal homologs showing increased abundance under sporulation-associated CCY conditions, including Cry32- and Cyt1-like proteins, were encoded on plasmid pBto-42_3, whereas the majority of degradative enzymes enriched in LB-derived extracellular fractions, including collagenases, chitinases, enhancin-like proteins, lipases and cytolysins, originated from chromosomal loci. Although further studies will be required to determine whether this distribution reflects coordinated regulatory mechanisms, the observed pattern suggests that plasmid-borne pesticidal factors and chromosomally encoded degradative enzymes may contribute differentially to the extracellular pathogenic repertoire of Bto_UNVM-42 under distinct physiological conditions.

The extracellular repertoire identified here includes multiple proteins previously associated with host invasion and tissue degradation in entomopathogenic bacteria. Collagenases, chitinases, and serine proteases may contribute to degradation of structural barriers and extracellular matrices, thereby facilitating tissue invasion and increasing host susceptibility (Fedhila et al., 2006; Geng et al., 2016). In addition, the detection of enhancin-like proteins further supports a cooperative pathogenic mechanism, since these metalloproteases are known to disrupt protective host matrices and enhance susceptibility to infection (Fang et al., 2009; Galloway et al., 2005). The coexistence of degradative enzymes with Bt-like pesticidal homologs may suggest a multifactorial extracellular strategy in which enzymatic degradation and membrane-associated toxicity may act synergistically. This interpretation is further supported by the nematode exposure assays, where concentrated cell-free supernatants induced severe loss of mobility that was not recovered after washing and progressive tissue disruption.

Importantly, the morphological alterations observed in dead *C. elegans* following exposure to concentrated extracellular fractions were consistent with those previously described for *P. redivivus* exposed to Bto_UNVM-42 cell-free supernatants (Sauka et al., 2026b). Although the two nematode species differ biologically, the similarity of the observed phenotypes supports the hypothesis that common extracellular factors may contribute to the nematicidal activity of this strain. Based on these and previous findings, we propose a multifactorial extracellular model of action (Figure 8) in which degradative enzymes may act in a coordinated manner. However, the specific contribution of each protein family remains unresolved. In particular, additional studies will be necessary to determine whether the detected Bt-like proteins are biologically active in the extracellular milieu, can interact with or be internalized by nematode tissues, and contribute directly to nematode mortality.

**Figure 8.**
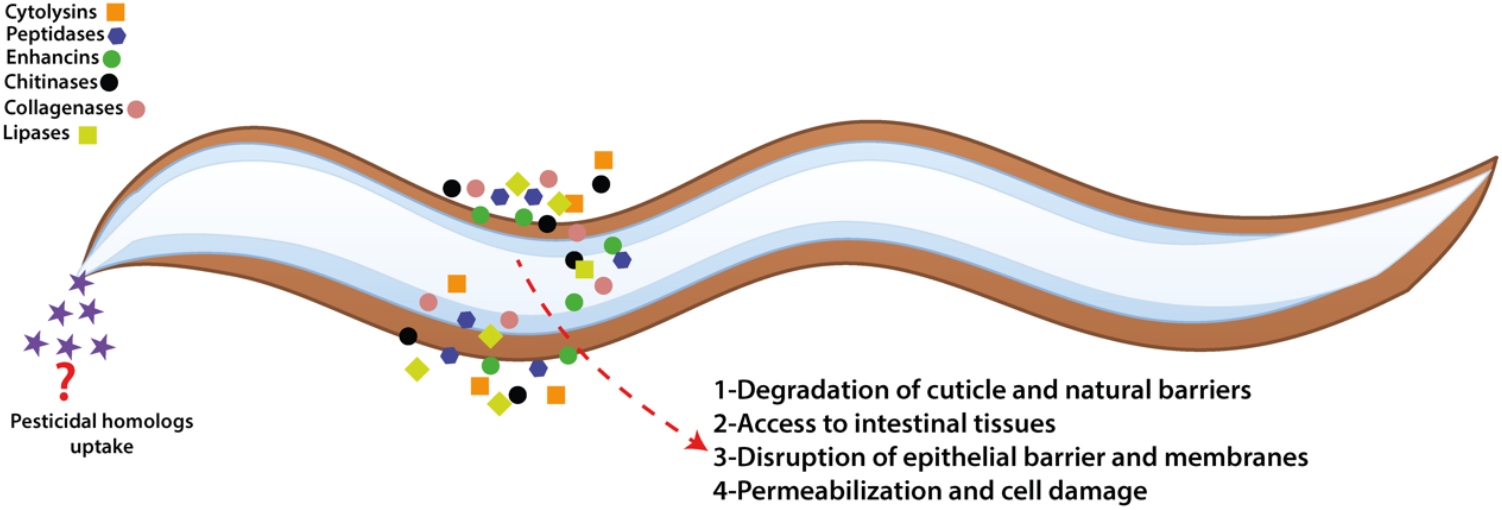
Proposed model of extracellular pathogenic mechanism in *B. toyonensis* Bto_UNVM-42 indicating different enzymes and the potential uptake of Bt-like pesticidal proteins.

Interestingly, the Cry32M-like homolog identified in the extracellular fraction, was previously detected in parasporal crystal-associated protein preparations of this strain (Sauka et al., 2026b). This observation suggests that certain Cry32-related proteins may occupy multiple physiological or spatial contexts, being associated not only with crystal preparations but also with soluble extracellular fractions. Together with the detection of Cyt1- and Mpp3-like proteins, these findings support a broader model in which Bt-related pesticidal proteins may occupy multiple spatial and physiological contexts beyond the classical crystal-associated paradigm.

Signal peptide analyses further indicated that extracellular localization may involve both classical and non-classical secretion pathways (Figure 4). While the Mpp3-like and Cyt1-like proteins displayed predicted N-terminal signal peptides consistent with classical secretion, the Cry32 K-, G-, M-, and U-like homologs lacked detectable signal peptides. Nevertheless, these Cry32-like homologs yielded positive SecretomeP prediction scores (SecP scores of 0.787, 0.916, 0.898, and 0.875, respectively), supporting the possibility of non-classical extracellular localization mechanisms. While several proteins displayed canonical secretion signals, the Cry32-like homologs and the S8 peptidase lacked predicted signal peptides but were identified as candidates for non-classical secretion. In *B. cereus* group bacteria, extracellular virulence-associated factors are frequently regulated by quorum-sensing and PlcR-associated pathways responsive to environmental conditions and nutrient availability (Gohar et al., 2008; Hajaij-Ellouze et al., 2006). Consequently, the extracellular repertoire observed here likely reflects a biologically regulated and condition-dependent pathogenic state.

Beyond its biological implications, this work also highlights the importance of incorporating proteomic profile analyses into the characterization of Bt and related bacteria isolates. Restricting characterization efforts exclusively to parasporal crystals may underestimate the functional diversity of these organisms, particularly when extracellular pesticidal homologs and virulence-associated enzymes are present. Integrating comparative extracellular proteomics with functional assays may therefore provide a more comprehensive understanding of Bt-related pathogenicity and ecological adaptation.

Overall, the results support a model in which *B. toyonensis* Bto_UNVM-42 deploys a flexible extracellular pathogenic repertoire composed of degradative enzymes, enhancin-like proteins, and Bt-related pesticidal homologs. This extracellular strategy extends beyond the traditional crystal-centric model of Bt pathogenicity and provides new insights into the molecular basis of nematicidal activity in *B. cereus* group bacteria.

## 5. Conclusion

This study provides comparative proteomic evidence that the extracellular fractions of *B. toyonensis* biovar thuringiensis Bto_UNVM-42 contain a diverse repertoire of proteins potentially associated with nematicidal activity, including Cry32-, Cyt1-, and Mpp3-like pesticidal homologs together with collagenases, chitinases, serine proteases, cytolysins, and enhancin-like proteins.

The consistent detection of these proteins in filtered cell-free supernatants indicates that biologically relevant Bt-related factors are present outside the classical spore–crystal complex. Comparative proteomic profiling analyses further revealed that extracellular protein composition is strongly influenced by growth conditions and physiological state, with sporulation-associated conditions favoring enrichment of Bt-like homologs and vegetative conditions favoring degradative extracellular enzymes.

Together with the observed nematicidal effects on *C. elegans*, these findings support a potential multifactorial, extracellular pathogenic model, in which degradative enzymes and membrane-active proteins may act cooperatively during host interaction and tissue disruption. Interestingly, comparative analyses also suggest a genomic partitioning of extracellular functions, with plasmid-encoded Bt-like pesticidal homologs predominating under sporulation-associated conditions and chromosomally encoded degradative enzymes being preferentially enriched during vegetative growth.

More broadly, this work expands the current understanding of Bt-related protein localization and highlights the importance of extracellular proteomics for the characterization of novel *B. cereus* group isolates. The results support an expanded model of Bt pathogenicity in which extracellular soluble factors contribute significantly to biological activity beyond the classical crystal-associated paradigm.

## CRediT authorship contribution statement

Conceptualization, L.P., methodology, S.R.-M., C.P and L.P.; formal analysis, S.R.-M., C.P, and L.P.; investigation, S.R.-M.; C.P. and L.P.; resources, L.P.; data curation, L.P.; writing-original draft preparation, L.P.; writing, review and editing, S.R.-M.; C.P. and L.P.; visualization, L.P.; supervision, L.P.; project administration, L.P.; funding acquisition, L.P. All authors have read and agreed to the published version of the manuscript.

## Declarations

### Ethical approval

This article does not contain any studies involving humans performed by any of the authors. Experiments involving nematodes were conducted in accordance with institutional and national guidelines applicable to invertebrate model organisms.

### Availability of data and materials

Supplementary data including the complete proteome of strain UNVM-42 and the raw LC–MS/MS proteomics datasets generated during this study are available at Zenodo under DOI: 10.5281/zenodo.19426829.

## Acknowledgements

Leopoldo Palma gratefully acknowledges the Spanish Ministry of Science, Innovation, and Universities, the Spanish State Research Agency, and the European Union for funding his Ramón y Cajal contract (grant ref. RYC2023-043507-I).

